# AniProtDB: A Collection of Uniformly Generated Metazoan Proteomes for Comparative Genomics Studies

**DOI:** 10.1101/2020.10.17.342964

**Authors:** Sofia N. Barreira, Anh-Dao Nguyen, Mark T. Fredriksen, Tyra G. Wolfsberg, R. Travis Moreland, Andreas D. Baxevanis

## Abstract

To address the void in the availability of high-quality proteomic data traversing the animal tree, we have implemented a pipeline for generating *de novo* assemblies based on publicly available data from the NCBI Sequence Read Archive, yielding a comprehensive collection of proteomes from 100 species spanning 21 animal phyla. We have also created the Animal Proteome Database (AniProtDB), a resource providing open access to this collection of high-quality metazoan proteomes, along with information on predicted proteins and protein domains for each taxonomic classification and the ability to perform sequence similarity searches against all proteomes generated using this pipeline. This solution vastly increases the utility of these data by removing the barrier to access for research groups who do not have the expertise or resources to generate these data themselves and enables the use of data from non-traditional research organisms that have the potential to address key questions in biomedicine.

**Database URL:** https://research.nhgri.nih.gov/aniprotdb

## Introduction

Comparative genomic analyses have provided keen insights into both the commonalities and differences between animal species, significantly advancing our understanding of genome structure, the processes underlying genome dynamics, the phylogenetic relationships between species, the evolution of gene families, population genomics, and the mechanisms underlying biological diversity, to name a few (Eisen and Fraser 2003; Hardison 2003; Elllegren 2008; Hardison and Taylor 2012; Lawrie and Petrov 2014). While the amount of sequence data contained within publicly available databases continues to increase at an exponential rate, encompassing 95% of all metazoan species, genomic data from invertebrate species continues to be quite underrepresented in these databases, with 91% of the data sets in the NCBI Sequence Read Archive (SRA) having been generated from vertebrate species alone (Kodama et al. 2012; Sayers et al. 2019a). This is not surprising, as there has been a traditional reliance on a small number of well-established vertebrate model organisms over the last several decades (Milinkovitch and Tzika 2007; Bolker 2014). Fortunately, there has recently been a renewed appreciation for the need to expand the number of tractable research organisms available to researchers (Russell et al. 2017; Sánchez Alvarado 2018; NIGMS/NIH 2019). Along with increased access to ever-improving sequencing technologies, this has resulted in numerous whole-genome sequencing efforts aimed at capturing and representing the breadth of diversity across the animal kingdom (GIGA Community of Scientists 2013), leading in turn to a corresponding exponential increase in the size of public data repositories such as GenBank (Sayers et al. 2019b; Baxevanis 2020).

While the availability of these myriad genomic, transcriptomic, and proteomic data sets provides a heretofore-unparalleled wealth of biological knowledge, it also poses a conundrum to most biologists in evaluating which data sets are of sufficient quality for use in designing, performing, and interpreting the results of their own experiments. Most publicly available data sets are comprised of raw sequencing reads that need to be processed, assembled, and annotated before meaningful information can be extracted from them, and while more and more researchers have the necessary skills to perform these tasks, access to the kinds of large-scale computational resources needed to perform these analyses on a wider scale still remains beyond the reach of many research groups. To address these concerns, we have developed and implemented a robust pipeline using freely available software components for generating *de novo* proteomes and associated protein domain information for representative species from as many animal phyla as possible, along with specific recommendations on how to effectively use these components. This pipeline is designed to foster reproducibility, increase the utility of publicly available sequence data, and provide a high degree of confidence when performing downstream comparative analyses. In addition, using data from SRA and availing ourselves of high-performance computing resources available to us, we have generated high-quality proteomes from 100 species spanning 21 animal phyla that are freely available through the Animal Proteome Database (AniProtDB). This new web-based resource removes the barriers to access to proteomic data from across the animal tree, providing open access to data that have been generated in a consistent fashion and with appropriate quality control measures, thereby enabling users to ultimately make confident biological conclusions.

### New Approaches

#### Proteome Generation Pipeline

The pipeline used to generate *de novo* proteomes based on transcriptome data found within SRA is illustrated in **Figure 1**. In the first of three checkpoints, FastQC provides multiple quality metrics for the RNA sequence reads, including the distribution of quality scores at each position as well as a list of over-represented sequences that represent more than 0.1% of total reads (indicating a significant degree of contamination). This output is used to set the parameters for Trim Galore! which trims sequencing adapters, over-represented sequences, and low-quality bases from the end of each read. The reads are then passed through a second FastQC checkpoint to ensure that the overall read quality has improved after the trimming step, applying the same gating criteria as before.

**Figure 1.**
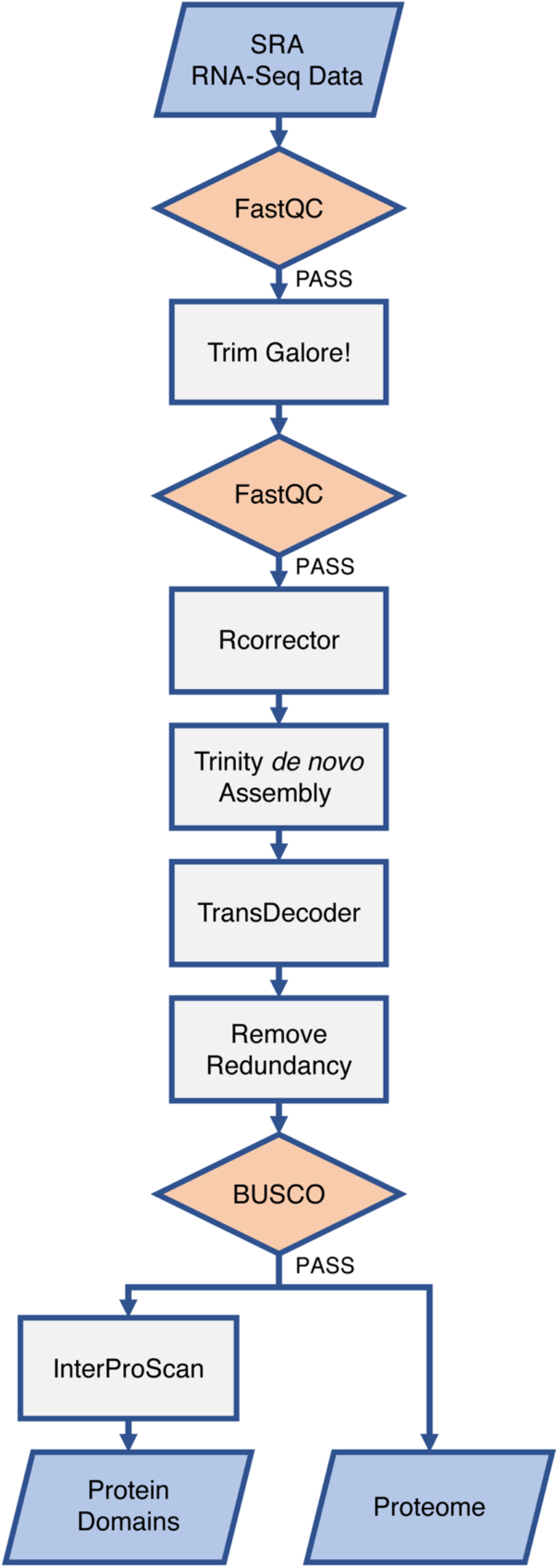
*De novo* metazoan proteome generation pipeline. The pipeline input (RNA-seq data from SRA) and output (Protein Domains and Proteomes) boxes are shaded in blue. Three quality control checkpoints (indicated by the orange-shaded boxes) are included and, if a data set scores below established thresholds at any step, it is immediately removed from further consideration (**Table S2**). See main text for details.

In the next series of steps, any random sequencing errors in the RNA-seq reads are corrected using Rcorrector (Song and Florea 2015) prior to *de novo* transcriptome assembly using Trinity (Haas et al. 2013). TransDecoder is then used to identify coding regions within the Trinity-generated transcript sequences. A custom script (see below) removes all peptides that are incomplete at the N- or C-terminal end, as well as any redundant proteins, ensuring that each protein sequence is represented once and only once so as to not skew statistics in the next step. We then use BUSCO (Simão et al. 2015; Waterhouse et al. 2017) to search this non-redundant proteome file for the presence of a comprehensive set of single-copy orthologs. As a third quality checkpoint, we have applied a BUSCO score threshold of 70% in order to retain proteomes from biologically interesting phyla that are not widely studied and to increase taxonomic breadth. In the final step, InterProScan is used to detect protein domains within each protein sequence. The pipeline yields two files as output: a FASTA-formatted file containing all of the protein sequences comprising the organism’s proteome (including transcript isoforms) and a text file with the InterProScan domain information from those sequences (**Table S1**). Detailed instructions for performing each step of the proteome pipeline (including scripts and usage examples) have been provided in the ‘Pipeline Overview’ section of the AniProtDB web site in order to facilitate reproducibility and the incorporation of future data sets using the same methodology.

#### The AniProtDB Database

The Animal Proteome DataBase (AniProtDB; https://research.nhgri.nih.gov/aniprotdb) houses the collection of metazoan proteomes generated by our pipeline. The ‘AniProtDB Statistics’ page contains informative metrics for each species represented in this database. A missing entry at the phylum or class level indicates that either there were no data available for that taxonomic group or the available data did not pass the quality control thresholds described above (**Table S2**). Predicted proteins and protein domain information for each phylum, class, and species classification can be found in the ‘Downloads’ section of the site, while the ‘Proteome Search’ and ‘Protein Domain Search’ tools can be used to interrogate the database for sequences and domains of interest. We have also implemented SequenceServer (Priyam et al. 2019), which allows users to perform sequence similarity searches of their data against either specific proteomes, combinations of proteomes, or all proteomes generated using our pipeline.

### Case study: An examination of the p53 protein domain across the Metazoa

Here, we provide a case study on interrogating proteomic data from AniProtDB to facilitate the analysis of the evolution of the p53 tumor suppressor gene, the origins of which can be traced back to the base of the animal tree (Belyi et al. 2010; Maxwell et al. 2014; Trigos et al. 2018). AniProtDB can be used to gather all of the proteins that possess the p53 domain across the animal kingdom by querying the ‘Protein Domain Search’ page using either the protein domain name (`P53`) or the CDD accession number (`cd08367`) in the search box (**Figure 2**). The results are returned in tabular format, showing the number of proteins in each species that contains the domain of interest. Users can also view the protein domain itself or the full-length domaincontaining protein. Of the 100 species in AniProtDB, 87 have at least one protein containing the p53 domain. For some species, AniProtDB also provides isoforms of the proteins containing this domain. With these data in hand, researchers can perform additional phylogenetic analyses and comparative biological studies with the relevant proteins from a multitude of species from different phyla and classes.

**Figure 2.**
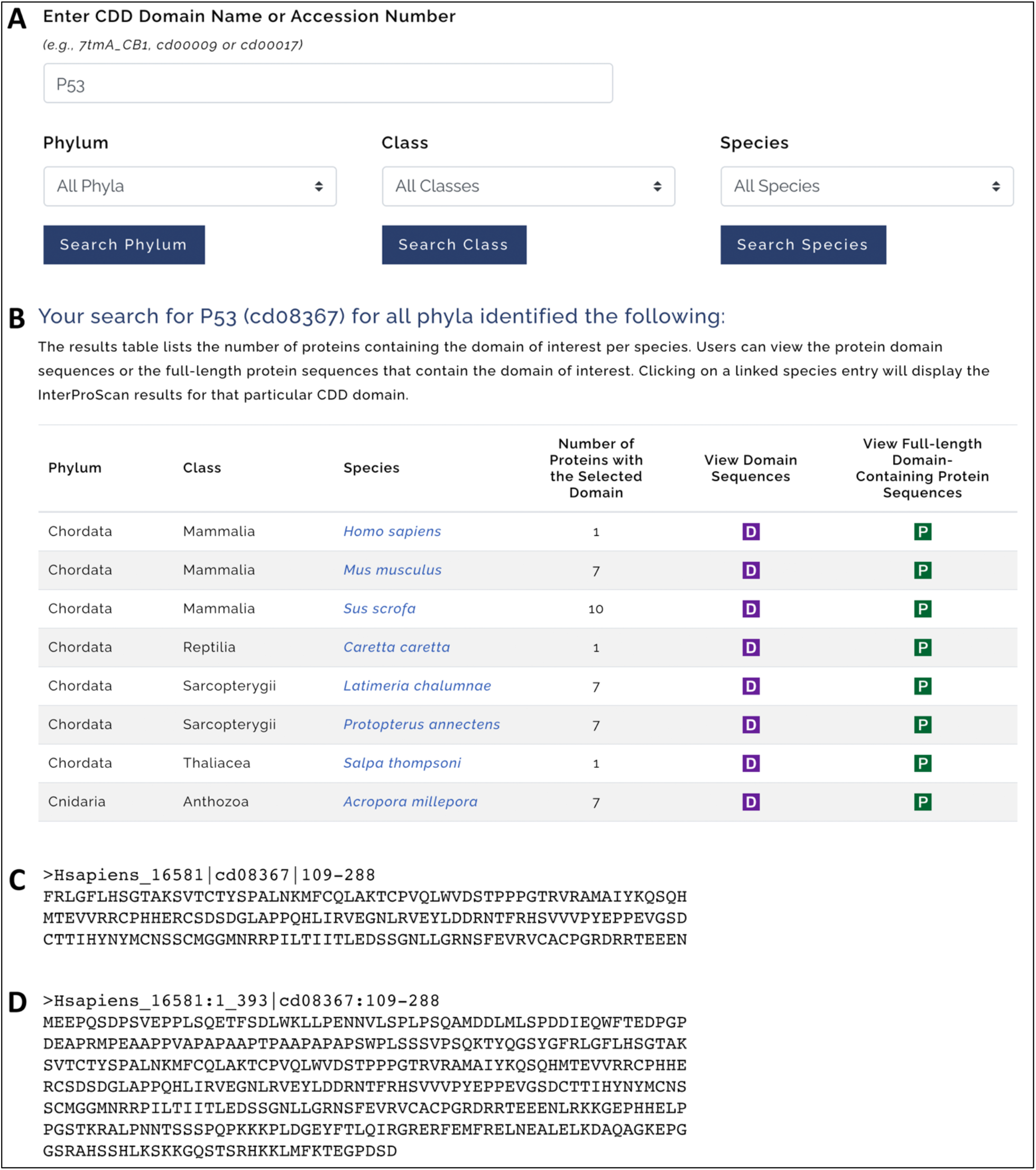
The protein domain search page in AniProtDB. (A) Users can enter a domain of interest in the search box (i.e., `P53` or CDD accession number `cd08367`). Results can be limited to a specific phylum, class, or species by selecting the available entries in the dropdown menus. (B) In the results table, here showing only human and the subsequent seven p53 entries for brevity, users can select the ‘D’ or ‘P’ buttons for each entry to view the truncated sequences of the domain or view the full-length proteins that contain the domain of interest, respectively. Figure sections C and D show an example of results for the *Homo sapiens* entry produced by clicking on the ‘D’ and ‘P’ buttons. The human proteome has one protein containing the p53 domain. Clicking the ‘D’ button will display the sequence of the domain only (C), extracted from the full-length protein (Hsapiens_16581), with the domain accession and coordinates in the sequence header (109-288). The ‘P’ button displays the full-length protein (D). A quick search on UniProt confirms that this protein is the human cellular tumor antigen p53 protein (UniProt accession P04637).

A further exploration of protein conservation across the Metazoa may be conducted using AniProtDB to perform a BLAST alignment search with either a DNA or protein sequence (**Figure 3**). Using the human p53 protein sequence shown in **Figure 2D** as the query against the proteomes of four emerging non-bilaterian research organisms (specifically, *Aurelia aurita, Hydra vulgaris, Hydractinia symbiolongicarpus*, and *Mnemiopsis leidyi*) returns the view shown in **Figure 3A**. (Alternatively, users may also perform a BLAST search against all proteome data contained within AniProtDB). The BLAST results include statistically significant alignments against AniProtDB proteins (**Figure 3B**), as well as links to additional information on protein domains and GO terms for each aligned AniProtDB protein sequence (**Figure 3C**). The BLAST results for the candidate p53 metazoan proteins indicate numerous high-scoring full-length candidates with very low E-values from all four of the non-bilaterian proteomes that were queried. Interestingly, BLAST detected the presence of the p53 protein domain in several statistically significant target sequences from the early branching invertebrate animal *Mnemiopsis leidyi* (Ryan et al., 2013), a notable result indicative of a high degree of p53 conservation from one end of the Metazoa to the other.

**Figure 3.**
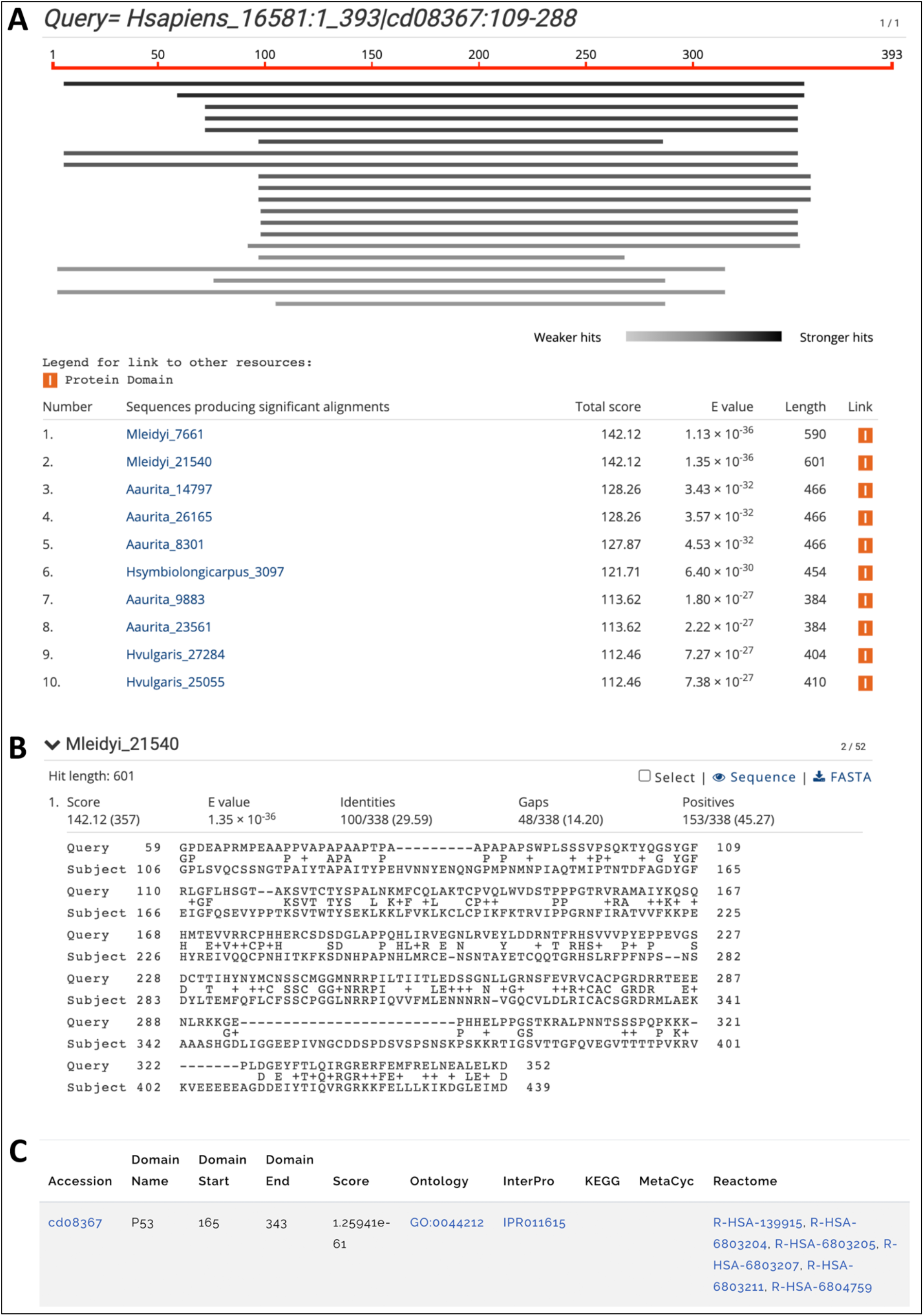
Example of alignment results from the BLAST section of AniProtDB. A BLASTP search was conducted using the sequence of the human protein p53 (Hsapiens_16581) as the query against the proteomic databases of four non-bilaterian invertebrate research organisms (*Aurelia aurita, Hydra vulgaris, Hydractinia symbiolongicarpus*, and *Mnemiopsis leidyi*). (A) The results page displays a graphic with the distribution of top hits and a summary table of hits found using BLASTP (with just the top ten hits displayed here for brevity). Clicking on any entry in the summary table will redirect users to the alignment visualization section. (B) Here, clicking on the longest and highest-scoring *Mnemiopsis* hit in the table takes the user to the alignment of the human p53 query with a protein from *M. leidyi* (Mleidyi_21540). (C) Clicking on the ‘I’ button on the right side of the results table (the ‘Link’ column in ‘A’) redirects users to the results of an InterProScan search of the *M. leidyi* protein against CDD. The links provide direct access to relevant entries in the GO, InterPro, KEGG, MetaCyc, and Reactome databases, respectively.

## Conclusions

In this study, we have implemented a pipeline for generating *de novo* assemblies of RNA-seq data to create a comprehensive collection of high-quality proteomes from representative metazoan species. The data sets derived from this pipeline directly address the under-representation and significant lack of data from numerous taxonomic groups by providing high-quality proteomes for at least one species from almost every class of metazoan phyla. These proteomes were all generated using consistent methodologies, quality control thresholds, and measures of completeness, and the adoption of a standardized pipeline such as the one described here will facilitate the generation of useful data sets for downstream analyses. We have made these proteomes freely available through the AniProtDB web site in order to both increase the utility of these data and remove the barrier to access for those who do not have the computational resources necessary to generate these data. This resource will be updated on a continual basis, with the community encouraged to provide suggestions or alert us to new data sets for inclusion. We foresee that this pipeline and resource will be of significant utility when applied to research projects across a wide spectrum of biological disciplines and further promote the adoption of promising new research organisms that are well-positioned for addressing key questions in biomedicine.

## Acknowledgments

This work utilized the Biowulf high-performance supercomputing resource of the Center for Information Technology at the National Institutes of Health (http://hpc.nih.gov).

