## Supplemental Tables 1 and 2 for "AniProtDB: A Collection of Uniformly Generated Metazoan Proteomes for Comparative Genomics Studies"

**Table S1.** Species included in AniProtDB

| Phylum | Class | Species | SRA Project | BUSCO Score | Proteome and Protein Domain Filenames |
|---|---|---|---|---|---|
| Annelida | Hirudinea | *Hirudinaria manillensis* | PRJNA548208 | 93.9 | Hmanillensis_proteins.fasta interproscan_Hmanillensis_CDD_results.txt |
| Annelida | Hirudinea | *Poecilobdella javanica* | PRJNA382185 | 77.0 | Pjavanica_proteins.fasta interproscan_Pjavanica_CDD_results.txt |
| Annelida | Oligochaeta | *Metaphire guillelmi* | PRJNA509281 | 86.7 | Mguillelmi_proteins.fasta interproscan_Mguillelmi_CDD_results.txt |
| Annelida | Oligochaeta | *Pheretima aspergillum* | PRJNA509281 | 85.8 | Paspergillum_proteins.fasta interproscan_Paspergillum_CDD_results.txt |
| Annelida | Polychaeta | *Riftia pachyptila* | PRJNA534438 | 86.5 | Rpachyptila_proteins.fasta interproscan_Rpachyptila_CDD_results.txt |
| Annelida | Polychaeta | *Urechis unicinctus* | PRJNA394029 | 86.3 | Uunicinctus_proteins.fasta interproscan_Uunicinctus_CDD_results.txt |
| Arthropoda | Arachnida | *Mesobuthus martensii* | PRJNA298268 | 91.1 | Mmartensii_proteins.fasta interproscan_Mmartensii_CDD_results.txt |
| Arthropoda | Arachnida | *Pardosa pseudoannulata* | PRJNA497774 | 95.2 | Ppseudoannulata_proteins.fasta interproscan_Ppseudoannulata_CDD_results.txt |
| Arthropoda | Branchiopoda | *Artemia franciscana* | PRJNA449186 | 75.3 | Afranciscana_proteins.fasta interproscan_Afranciscana_CDD_results.txt |
| Arthropoda | Branchiopoda | *Daphnia magna* | PRJNA533017 | 87.3 | Dmagna_proteins.fasta interproscan_Dmagna_CDD_results.txt |
| Arthropoda | Chilopoda | *Scolopocryptops sexspinosus* | PRJNA340270 | 73.1 | Ssexspinosus_proteins.fasta interproscan_Ssexspinosus_CDD_results.txt |
| Arthropoda | Chilopoda | *Strigamia maritima* | PRJNA386859 | 89.2 | Smaritima_proteins.fasta interproscan_Smaritima_CDD_results.txt |
| Arthropoda | Diplopoda | *Andrognathus corticarius* | PRJNA434333 | 71.8 | Acorticarius_proteins.fasta interproscan_Acorticarius_CDD_results.txt |
| Arthropoda | Diplopoda | *Californiulus sp.* | PRJNA434333 | 73.1 | Californiulussp_proteins.fasta interproscan_Californiulussp_CDD_results.txt |
| Arthropoda | Entognatha | *Holacanthella duospinosa* | PRJNA384703 | 84.9 | Hduospinosa_proteins.fasta interproscan_Hduospinosa_CDD_results.txt |
| Arthropoda | Entognatha | *Sinella curviseta* | PRJNA494047 | 83.1 | Scurviseta_proteins.fasta interproscan_Scurviseta_CDD_results.txt |
| Arthropoda | Insecta | *Apis mellifera* | PRJNA510543 | 91.2 | Amellifera_proteins.fasta interproscan_Amellifera_CDD_results.txt |
| Arthropoda | Insecta | *Spodoptera litura* | PRJNA511360 | 91.8 | Slitura_proteins.fasta interproscan_Slitura_CDD_results.txt |
| Arthropoda | Malacostraca | *Carcinus maenas* | PRJNA400568 | 87.1 | Cmaenas_proteins.fasta interproscan_Cmaenas_CDD_results.txt |
| Arthropoda | Malacostraca | *Penaeus vannamei* | PRJNA495361 | 80.2 | Pvannamei_proteins.fasta interproscan_Pvannamei_CDD_results.txt |
| Arthropoda | Maxillopoda | *Neolepas marisindica* | PRJNA516093 | 87.5 | Nmarisindica_proteins.fasta interproscan_Nmarisindica_CDD_results.txt |
| Arthropoda | Maxillopoda | *Tigriopus californicus* | PRJNA504307 | 82.6 | Tcalifornicus_proteins.fasta interproscan_Tcalifornicus_CDD_results.txt |

| | | | | | |
|---|---|---|---|---|---|
| Arthropoda | Merostomata | *Tachypleus tridentatus* | PRJNA483308 | 92.7 | `Ttridentatus_proteins.fasta` `interproscan_Ttridentatus_CDD_results.txt` |
| Brachiopoda | Craniforma | *Novocrania anomala* | PRJNA295688 | 74.2 | `Nanomala_proteins.fasta` `interproscan_Nanomala_CDD_results.txt` |
| Brachiopoda | Lingulata | *Lingula anatina* | PRJNA286275 | 75.6 | `Lanatina_proteins.fasta` `interproscan_Lanatina_CDD_results.txt` |
| Brachiopoda | Rhynconellata | *Liothyrella uva* | PRJNA314406 | 75.2 | `Luva_proteins.fasta` `interproscan_Luva_CDD_results.txt` |
| Brachiopoda | Rhynconellata | *Terebratalia transversa* | PRJNA295688 | 73.4 | `Ttransversa_proteins.fasta` `interproscan_Ttransversa_CDD_results.txt` |
| Bryozoa | Gymnolaemata | *Bugula neritina* | PRJNA244120 | 84.5 | `Bneritina_proteins.fasta` `interproscan_Bneritina_CDD_results.txt` |
| Bryozoa | Gymnolaemata | *Membranipora membranacea* | PRJNA289347 | 82.5 | `Mmembranacea_proteins.fasta` `interproscan_Mmembranacea_CDD_results.txt` |
| Chaetognatha | Sagittoidea | *Paraspadella gotoi* | PRJNA487918 | 71.0 | `Pgotoi_proteins.fasta` `interproscan_Pgotoi_CDD_results.txt` |
| Chaetognatha | Sagittoidea | *Sagitta elegans* | PRJNA487918 | 73.9 | `Selegans_proteins.fasta` `interproscan_Selegans_CDD_results.txt` |
| Chordata | Actinopterygii | *Acipenser baerii* | PRJNA274436 | 94.7 | `Abaerii_proteins.fasta` `interproscan_Abaerii_CDD_results.txt` |
| Chordata | Actinopterygii | *Thunnus maccoyii* | PRJNA288188 | 88.9 | `Tmaccoyii_proteins.fasta` `interproscan_Tmaccoyii_CDD_results.txt` |
| Chordata | Agnatha | *Eptatretus burgeri* | PRJNA371391 | 90.4 | `Eburgeri_proteins.fasta` `interproscan_Eburgeri_CDD_results.txt` |
| Chordata | Agnatha | *Lethenteron japonicum* | PRJNA371391 | 76.5 | `Ljaponicum_proteins.fasta` `interproscan_Ljaponicum_CDD_results.txt` |
| Chordata | Amphibia | *Rana temporaria* | PRJNA503663 | 95.4 | `Rtemporaria_proteins.fasta` `interproscan_Rtemporaria_CDD_results.txt` |
| Chordata | Amphibia | *Rhinatrema bivittatum* | PRJEB25429 | 94.6 | `Rbivittatum_proteins.fasta` `interproscan_Rbivittatum_CDD_results.txt` |
| Chordata | Ascidiacea | *Ciona intestinalis* | PRJNA248170 | 73.7 | `Cintestinalis_proteins.fasta` `interproscan_Cintestinalis_CDD_results.txt` |
| Chordata | Aves | *Anas platyrhynchos* | PRJNA563502 | 77.8 | `Aplatyrhynchos_proteins.fasta` `interproscan_Aplatyrhynchos_CDD_results.txt` |
| Chordata | Chondrichthyes | *Leucoraja erinacea* | PRJNA288370 | 78.1 | `Lerinacea_proteins.fasta` `interproscan_Lerinacea_CDD_results.txt` |
| Chordata | Chondrichthyes | *Okamejei kenojei* | PRJNA438719 | 87.9 | `Okenojei_proteins.fasta` `interproscan_Okenojei_CDD_results.txt` |
| Chordata | Chondrichthyes | *Scyliorhinus torazame* | PRJNA371391 | 90.3 | `Storazame_proteins.fasta` `interproscan_Storazame_CDD_results.txt` |
| Chordata | Leptocardii | *Branchiostoma lanceolatum* | PRJNA416866 | 72.1 | `Blanceolatum_proteins.fasta` `interproscan_Blanceolatum_CDD_results.txt` |
| Chordata | Mammalia | *Homo sapiens* | PRJNA288500 | 86.0 | `Hsapiens_proteins.fasta` `interproscan_Hsapiens_CDD_results.txt` |
| Chordata | Mammalia | *Mus musculus* | PRJNA288500 | 83.1 | `Mmusculus_proteins.fasta` `interproscan_Mmusculus_CDD_results.txt` |
| Chordata | Mammalia | *Sus scrofa* | PRJNA481305 | 73.9 | `Sscrofa_proteins.fasta` `interproscan_Sscrofa_CDD_results.txt` |

| | | | | | |
|---|---|---|---|---|---|
| Chordata | Reptilia | *Caretta caretta* | PRJNA663187 | 92.6 | Ccaretta_proteins.fasta<br>interproscan_Ccaretta_CDD_results.txt |
| Chordata | Sarcopterygii | *Latimeria chalumnae* | PRJDB1984 | 85.3 | Lchalumnae_proteins.fasta<br>interproscan_Lchalumnae_CDD_results.txt |
| Chordata | Sarcopterygii | *Protopterus annectens* | PRJNA282925 | 88.0 | Pannectens_proteins.fasta<br>interproscan_Pannectens_CDD_results.txt |
| Chordata | Thaliacea | *Salpa thompsoni* | PRJNA279245 | 83.9 | Sthompsoni_proteins.fasta<br>interproscan_Sthompsoni_CDD_results.txt |
| Cnidaria | Anthozoa | *Acropora millepora* | PRJNA473876 | 82.6 | Amillepora_proteins.fasta<br>interproscan_Amillepora_CDD_results.txt |
| Cnidaria | Anthozoa | *Exaiptasia pallida* | PRJDB7145 | 80.0 | Epallida_proteins.fasta<br>interproscan_Epallida_CDD_results.txt |
| Cnidaria | Cubozoa | *Alatina alata* | PRJNA312373 | 77.3 | Aalata_proteins.fasta<br>interproscan_Aalata_CDD_results.txt |
| Cnidaria | Cubozoa | *Tripedalia cystophora* | PRJNA498176 | 87.8 | Tcystophora_proteins.fasta<br>interproscan_Tcystophora_CDD_results.txt |
| Cnidaria | Hydrozoa | *Hydra vulgaris* | PRJNA497966 | 89.6 | Hvulgaris_proteins.fasta<br>interproscan_Hvulgaris_CDD_results.txt |
| Cnidaria | Hydrozoa | *Hydractinia symbiolongicarpus* | PRJNA237004 | 87.4 | Hsymbiolongicarpus_proteins.fasta<br>interproscan_Hsymbiolongicarpus_CDD_results.txt |
| Cnidaria | Scyphozoa | *Aurelia aurita* | PRJNA252562 | 74.6 | Aaurita_proteins.fasta<br>interproscan_Aaurita_CDD_results.txt |
| Cnidaria | Scyphozoa | *Nemopilema nomurai* | PRJNA415234 | 80.5 | Nnomurai_proteins.fasta<br>interproscan_Nnomurai_CDD_results.txt |
| Ctenophora | Tentaculata | *Bolinopsis mikado* | PRJDB8655 | 77.6 | Bmikado_proteins.fasta<br>interproscan_Bmikado_CDD_results.txt |
| Ctenophora | Tentaculata | *Mnemiopsis leidyi* | PRJEB28334 | 83.0 | Mleidyi_proteins.fasta<br>interproscan_Mleidyi_CDD_results.txt |
| Echinodermata | Asteroidea | *Patiria pectinifera* | PRJNA371229 | 78.8 | Ppectinifera_proteins.fasta<br>interproscan_Ppectinifera_CDD_results.txt |
| Echinodermata | Asteroidea | *Pisaster ochraceus* | PRJNA603587 | 84.2 | Pochraceus_proteins.fasta<br>interproscan_Pochraceus_CDD_results.txt |
| Echinodermata | Crinoidea | *Antedon mediterranea* | PRJNA432136 | 74.9 | Amediterranea_proteins.fasta<br>interproscan_Amediterranea_CDD_results.txt |
| Echinodermata | Echinoidea | *Mesocentrotus franciscanus* | PRJNA531463 | 88.8 | Mfranciscanus_proteins.fasta<br>interproscan_Mfranciscanus_CDD_results.txt |
| Echinodermata | Echinoidea | *Paracentrotus lividus* | PRJNA392084 | 77.3 | Plividus_proteins.fasta<br>interproscan_Plividus_CDD_results.txt |
| Echinodermata | Holothuroidea | *Apostichopus japonicus* | PRJNA597445 | 82.3 | Ajaponicus_proteins.fasta<br>interproscan_Ajaponicus_CDD_results.txt |
| Echinodermata | Holothuroidea | *Apostichopus parvimensis* | PRJNA296688 | 88.2 | Aparvimensis_proteins.fasta<br>interproscan_Aparvimensis_CDD_results.txt |
| Echinodermata | Ophiuroidea | *Amphipholis kochii* | PRJDB8281 | 84.6 | Akochii_proteins.fasta<br>interproscan_Akochii_CDD_results.txt |
| Echinodermata | Ophiuroidea | *Amphiura filiformis* | PRJNA349786 | 86.5 | Afiliformis_proteins.fasta<br>interproscan_Afiliformis_CDD_results.txt |
| Hemichordata | Enteropneusta | *Ptychodera flava* | PRJNA284280 | 91.7 | Pflava_proteins.fasta<br>interproscan_Pflava_CDD_results.txt |

| | | | | | |
|---|---|---|---|---|---|
| Hemichordata | Enteropneusta | *Schizocardium californicum* | PRJNA296693 | 74.7 | Scalifornicum_proteins.fasta interproscan_Scalifornicum_CDD_results.txt |
| Micrognathozoa | Micrognathozoa | *Limnognathia maerski* | PRJNA289337 | 75.1 | Lmaerski_proteins.fasta interproscan_Lmaerski_CDD_results.txt |
| Mollusca | Aplacophora | *Stylomenia sulcodoryata* | PRJNA445319 | 80.3 | Ssulcodoryata_proteins.fasta interproscan_Ssulcodoryata_CDD_results.txt |
| Mollusca | Bivalvia | *Crassostrea gigas* | PRJNA543621 | 83.3 | Cgigas_proteins.fasta interproscan_Cgigas_CDD_results.txt |
| Mollusca | Bivalvia | *Mytilus edulis* | PRJNA525607 | 84.3 | Medulis_proteins.fasta interproscan_Medulis_CDD_results.txt |
| Mollusca | Cephalopoda | *Octopus bimaculoides* | PRJNA285380 | 91.3 | Obimaculoides_proteins.fasta interproscan_Obimaculoides_CDD_results.txt |
| Mollusca | Cephalopoda | *Sepia esculenta* | PRJNA471792 | 92.8 | Sesculenta_proteins.fasta interproscan_Sesculenta_CDD_results.txt |
| Mollusca | Gastropoda | *Ambigolimax valentianus* | PRJDB3972 | 81.4 | Avalentianus_proteins.fasta interproscan_Avalentianus_CDD_results.txt |
| Mollusca | Gastropoda | *Pomacea canaliculata* | PRJNA476647 | 85.9 | Pcanaliculata_proteins.fasta interproscan_Pcanaliculata_CDD_results.txt |
| Nematoda | Chromadorea | *Brugia malayi* | PRJNA294263 | 95.1 | Bmalayi_proteins.fasta interproscan_Bmalayi_CDD_results.txt |
| Nematoda | Enoplea | *Enoplus brevis* | PRJEB7588 | 75.0 | Ebrevis_proteins.fasta interproscan_Ebrevis_CDD_results.txt |
| Nematoda | Enoplea | *Trichinella pseudospiralis* | PRJNA450330 | 74.4 | Tpseudospiralis_proteins.fasta interproscan_Tpseudospiralis_CDD_results.txt |
| Nemertea | Hoplonemertea | *Antarctonemertes valida* | PRJNA485632 | 70.4 | Avalida_proteins.fasta interproscan_Avalida_CDD_results.txt |
| Nemertea | Pilidiophora | *Lineus sanguineus* | PRJNA322119 | 82.9 | Lsanguineus_proteins.fasta interproscan_Lsanguineus_CDD_results.txt |
| Nemertea | Pilidiophora | *Ramphogordius lacteus* | PRJNA322119 | 76.3 | Rlacteus_proteins.fasta interproscan_Rlacteus_CDD_results.txt |
| Phoronida | See note 1 | *Phoronis australis* | PRJNA393252 | 82.0 | Paustralis_proteins.fasta interproscan_Paustralis_CDD_results.txt |
| Placozoa | See note 1 | *Hoilungia hongkongensis* | PRJNA377631 | 89.5 | Hhongkongensis_proteins.fasta interproscan_Hhongkongensis_CDD_results.txt |
| Placozoa | See note 1 | *Trichoplax adhaerens* | PRJNA394204 | 79.6 | Tadhaerens_proteins.fasta interproscan_Tadhaerens_CDD_results.txt |
| Platyhelminthes | Catenulida | *Stenostomum sthenum* | PRJNA275074 | 70.2 | Ssthenum_proteins.fasta interproscan_Ssthenum_CDD_results.txt |
| Platyhelminthes | Rhabditophora | *Girardia guanajuatiensis* | PRJNA551297 | 74.6 | Gguanajuatiensis_proteins.fasta interproscan_Gguanajuatiensis_CDD_results.txt |
| Platyhelminthes | Rhabditophora | *Schmidtea mediterranea* | PRJNA390190 | 76.5 | Smediterranea_proteins.fasta interproscan_Smediterranea_CDD_results.txt |
| Porifera | Calcarea | *Grantia compressa* | PRJNA316185 | 71.0 | Gcompressa_proteins.fasta interproscan_Gcompressa_CDD_results.txt |
| Porifera | Calcarea | *Sycon ciliatum* | PRJEB7138 | 80.0 | Sciliatum_proteins.fasta interproscan_Sciliatum_CDD_results.txt |
| Porifera | Demospongiae | *Halichondria panicea* | PRJNA594151 | 85.6 | Hpanicea_proteins.fasta interproscan_Hpanicea_CDD_results.txt |

| | | | | | |
|---|---|---|---|---|---|
| Porifera | Homoscleromorpha | *Corticium candelabrum* | PRJNA162903 | 78.0 | `Ccandelabrum_proteins.fasta` `interproscan_Ccandelabrum_CDD_results.txt` |
| Priapulida | Halicryptomorpha | *Halicryptus spinulosus* | PRJNA295688 | 88.6 | `Hspinulosus_proteins.fasta` `interproscan_Hspinulosus_CDD_results.txt` |
| Rotifera | Monogononta | *Brachionus koreanus* | PRJNA267756 | 79.5 | `Bkoreanus_proteins.fasta` `interproscan_Bkoreanus_CDD_results.txt` |
| Rotifera | Monogononta | *Brachionus manjavacas* | PRJNA345262 | 77.7 | `Bmanjavacas_proteins.fasta` `interproscan_Bmanjavacas_CDD_results.txt` |
| Tardigrada | Eutardigrada | *Hypsibius dujardini* | PRJDB5599 | 76.3 | `Hdujardini_proteins.fasta` `interproscan_Hdujardini_CDD_results.txt` |
| Tardigrada | Eutardigrada | *Paramacrobiotus richtersi* | PRJNA369152 | 78.9 | `Prichtersi_proteins.fasta` `interproscan_Prichtersi_CDD_results.txt` |

1. No class names assigned in this phylum.

**Table S2.** Number of SRA data sets that did not meet quality control thresholds, by phylum

| Phylum | Total Failed Sets | First FastQC Step | Second FastQC Step | BUSCO Step |
|---|---|---|---|---|
| Annelida | 2 | 0 | 0 | 2 |
| Arthropoda | 29 | 10 | 7 | 12 |
| Brachiopoda | 1 | 0 | 0 | 1 |
| Bryozoa | 2 | 0 | 2 | 0 |
| Chordata | 9 | 3 | 0 | 6 |
| Cnidaria | 5 | 3 | 2 | 0 |
| Ctenophora | 6 | 0 | 1 | 5 |
| Echinodermata | 14 | 0 | 3 | 11 |
| Entoprocta | 3 | 1 | 0 | 2 |
| Gastrotricha | 3 | 1 | 0 | 2 |
| Gnathostomulida | 2 | 0 | 0 | 2 |
| Hemichordata | 3 | 1 | 1 | 1 |
| Mollusca | 15 | 0 | 1 | 14 |
| Nematoda | 23 | 5 | 0 | 18 |
| Nemertea | 5 | 0 | 1 | 4 |
| Onychophora | 3 | 0 | 0 | 3 |
| Phoronida | 3 | 0 | 0 | 3 |
| Platyhelminthes | 6 | 2 | 1 | 3 |
| Porifera | 13 | 4 | 2 | 7 |
| Priapulida | 2 | 0 | 0 | 2 |
| Rotifera | 4 | 0 | 0 | 4 |
| Tardigrada | 3 | 0 | 0 | 3 |
| ***Total*** | 156 | 30 | 21 | 105 |

The *Total Failed Sets* column indicates the total number of data sets that were rejected at any of the checkpoints in the proteome generation pipeline. The rightmost three columns indicate the number of data sets that failed at each of the three checkpoints.
